# Dynamic alterations of dural and bone marrow B cells in an animal model of progressive multiple sclerosis

**DOI:** 10.1101/2024.08.23.609437

**Authors:** Alexandra Florescu, Michelle Zuo, Angela A. Wang, Kevin Champagne-Jorgensen, Mohammed Ariyan Noor, Lesley A. Ward, Erwin van Puijenbroek, Christian Klein, Jennifer L. Gommerman

## Abstract

In multiple sclerosis (MS), the leptomeninges (LM) are populated with immune cell aggregates that correlate with disease progression. The impact of LM inflammation on the adjacent dura is largely unknown. Using a mouse model of MS that induces brain LM inflammation and age-dependent disease progression, we found that encephalitogenic T cells and B220^high^ B cells accumulate substantially in the brain LM and parenchyma of both young and aged mice, while the adjacent dura remains relatively inert. We also observed a population of anti-CD20 resistant B220^low^ B cells in the dura and bone marrow that virtually disappear at disease onset and accumulate in the brain of young mice concomitant with disease remission. In contrast, aged mice show a paucity of brain-resident B220^low^ B cells at the expense of class-switched B220^high^ B cells concomitant with severe, chronic disease. In summary, dynamic changes in brain, LM and dural B cells are associated with age-dependent disease severity in an animal model of progressive MS.

**Short Summary:** Florescu *et al*. investigate the temporal accumulation of immune cells within distinct meningeal compartments in an animal model of progressive MS and uncover a population of anti-CD20 resistant dural B cells that remain in the brain parenchyma at disease remission.

## Introduction

The meninges, an anatomical buffer that has evolved to protect the central nervous system (CNS) from pathogens, are comprised of three distinct layers: the dura mater found immediately underneath the bony skull, the arachnoid mater which forms the space through which cerebrospinal fluid (CSF) flows, and the pia mater which overlays the brain parenchyma. The arachnoid mater and pia mater are collectively termed the leptomeninges (LM). Whereas the LM is lymphocyte-poor, the dura mater contains a variety of immune cells including T cells, neutrophils, mature B cells and developing B cells (Rustenhoven et al., 2021, Brioschi et al., 2021, Schafflick et al., 2021, Wang et al., 2021b, Rua and McGavern, 2018).

The immune cell-permissive state of the dura is largely due to fenestrated and innervated blood vessels that are more permeable than those in the brain or LM, dural lymphatic drainage of the brain and CSF, and osseous channels that travel from the skull bone marrow (BM) to the dura. The latter provide a route for BM cells, such as developing B cells, to enter the dura (Brioschi et al., 2021, Cugurra et al., 2021). The involvement of the dura as a source of immune cells in CNS autoimmune inflammation, such as that seen in multiple sclerosis (MS), remains unclear.

MS is an autoimmune disease of the brain and spinal cord in which immune cell subsets collaborate to cause damage to the CNS, including demyelination and axonal loss. Younger patients in early stages of the disease typically present with relapsing remitting MS (RRMS), which is characterized by transient episodes of clinical worsening followed by partial or full recovery (Lublin and Reingold, 1996). Clinical presentation of the disease changes with age, with individuals exhibiting increasing disability without recovery (Lublin and Reingold, 1996, Scalfari et al., 2011). This phase of the disease has been called progressive MS (PMS). The age-dependent mechanisms driving MS progression are not well understood.

Pathologically, focal demyelinating lesions in the white matter (WM) are the hallmark of MS. While new WM lesions are observed in relapsing MS, they are less common in PMS. Instead, areas of grey matter (GM) demyelination directly under the pia (subpial lesions) have been associated with cortical atrophy and disease progression, particularly in aged patients. Despite evidence of axonal and neuronal loss within demyelinated GM lesions, GM lesions are relatively lymphocyte and complement poor compared to WM lesions (van Horssen et al., 2007, Brink et al., 2005, Peterson et al., 2001, Lucchinetti et al., 2011). While the etiology of these GM lesions is debated, one leading hypothesis is that compartmentalized inflammation in the adjacent LM drives GM damage through soluble factors produced by LM-resident immune cells that cross through the glial limitans superficialis into the cortex where they can have injurious properties (Magliozzi et al., 2007).

The LM are an important site of pathogenic T cell entry during the onset of experimental autoimmune encephalomyelitis (EAE) (Lodygin et al., 2019, Lodygin et al., 2013, Schlager et al., 2016). We previously showed that the LM of SJL/J mice undergo remarkable stromal cell remodeling in response to adoptive transfer of Th17 cells, enabling the formation of a niche that supports further immune cell influx and Th17 cell maintenance and polarization (Pikor et al., 2015). These LM aggregates are adjacent to regions of cortical pathology, including GM demyelination, microglial activation and glia limitans disruption, as observed in MS (Pikor et al., 2015, Ward et al., 2020). Moreover, adoptive transfer SJL/J EAE mice exhibit age-dependent disease severity and chronicity. While young mice clinically remit and resolve LM inflammation and cortical pathology, aged mice develop persistent clinical disease, LM inflammation and cortical pathology as well as evidence of neurodegeneration including axonal stress, synapse loss and cortical thinning (Zuo et al., 2022). Testing of MS therapies in the adoptive transfer SJL/J EAE model has also positioned the LM as an important site of drug action. Indeed, we recently showed that depletion of B cells with anti-CD20 in SJL/J mice reduces the size of LM aggregates and prevents GM demyelination and cortical injury (Wang et al., 2024).

The role of the dura in MS and EAE has only been recently studied. Reports using C57BL/6 active EAE have described an increase in dura-resident immune cells including myeloid cells, IgG^+^ and IgM^+^ plasma cells (PCs), Th17 and regulatory T cells, while IgA^+^ PCs were decreased compared to naïve mice (Louveau et al., 2018, Rustenhoven et al., 2021, Schafflick et al., 2021, Cugurra et al., 2021). Ablation of dural lymphatics prior to EAE induction delayed disease onset and decreased disease severity, possibly due to reduced antigen and immune cell drainage to sites of activation such as areas adjacent to dural sinuses or the cervical lymph nodes (LNs) (Louveau et al., 2018). Interestingly, neutrophils, Gr-1^+^ myeloid cells and T cells have been found to move from the dura to the LM during EAE through arachnoid cuff exit points which connect the dura and LM, suggesting that the dura may be a source of potentially injurious immune cells (Smyth et al., 2024). Taken together, data from C57BL/6 EAE supports a role for the dura as a potential source of immune cells that invade the LM and parenchyma and/or as a conduit for antigen and immune cells out of the LM and parenchyma. However, other studies have found no difference in EAE course or immune cell infiltration into the CNS, LM or dura following dural lymphatic ablation (Merlini et al., 2022, Li et al., 2023). Moreover, C57BL/6 EAE mice do not exhibit immune cell aggregates in the brain LM, nor do they show evidence of subpial cortical GM pathology, limiting our understanding of immune cell movement between the brain dura and the inflamed LM during EAE. Indeed, in a rat model of GM damage, the relative accumulation of T cells and myeloid cells in the dura was significantly lower than that seen in the LM or brain of the same rats (Merlini et al., 2022).

To better characterize the temporal accumulation of different types of immune cells in distinct layers of the brain meninges in the context of brain neuroinflammation, we applied our model of EAE induced by adoptive transfer of encephalitogenic proteolipid protein (PLP)-primed Th17 cells into both young and aged SJL/J mice. We probed the role of age on the accumulation of immune cells in the dura, LM, brain, skull BM and femur BM at multiple timepoints of the EAE disease trajectory. Independent of age, we found that compared to the LM and the brain parenchyma, the dura is not a site for accumulation of encephalitogenic Th17 cells. However, we noted that developing/immature B220^low^ B cells disappeared from the dura and BM during EAE, and that different subsets of B cells, some of which resist anti-CD20 therapy, exhibited an age-dependent brain compartmentalization at disease remission.

## Results and discussion

### Adoptively transferred encephalitogenic Th17 cells are recovered from the leptomeninges and brain, but not the dura, during EAE

To evaluate the relative accumulation of encephalitogenic Th17 cells in a setting with sub-pial cortical GM injury, we tracked pathogenic T cell infiltration into the dura, LM and brain by adoptive transfer of CFSE labelled Th17-primed PLP-reactive cells into recipient SJL/J mice. Similar to MS, this model induces stromal cell remodeling and the formation of lymphocyte aggregates that contain discrete T cell and B cell zones adjacent to areas of GM demyelination, microglial activation and glial limitans disruption (Pikor et al., 2015, Ward et al., 2020).

Accordingly, pre-primed encephalitogenic Th17 cells were labelled with CFSE then transferred into young (6-to 10-week-old) recipient mice. Single cell suspensions of the dura, LM, brain, blood and spleen were analyzed by flow cytometry at several timepoints following adoptive transfer (Fig. 1A, Supp. Fig. 1A). During the pre-onset phase (days 1-5 following adoptive transfer), CFSE^+^ T cells were detected in the blood and spleen as early as day 1 but not in either the dura or LM (Fig. 1B). At disease onset (day 4 onwards), CFSE^+^ T cells were observed in the spleen from day 4 to 7, but by day 11 were no longer detectable. As early as day 5, CFSE^+^ T cells were observed in the LM and brain of mice and continued to accumulate up until peak disease at day 11. In contrast, no CFSE^+^ T cell infiltration occurred in the dura at any of these timepoints (Fig. 1C).

**Fig. 1:**
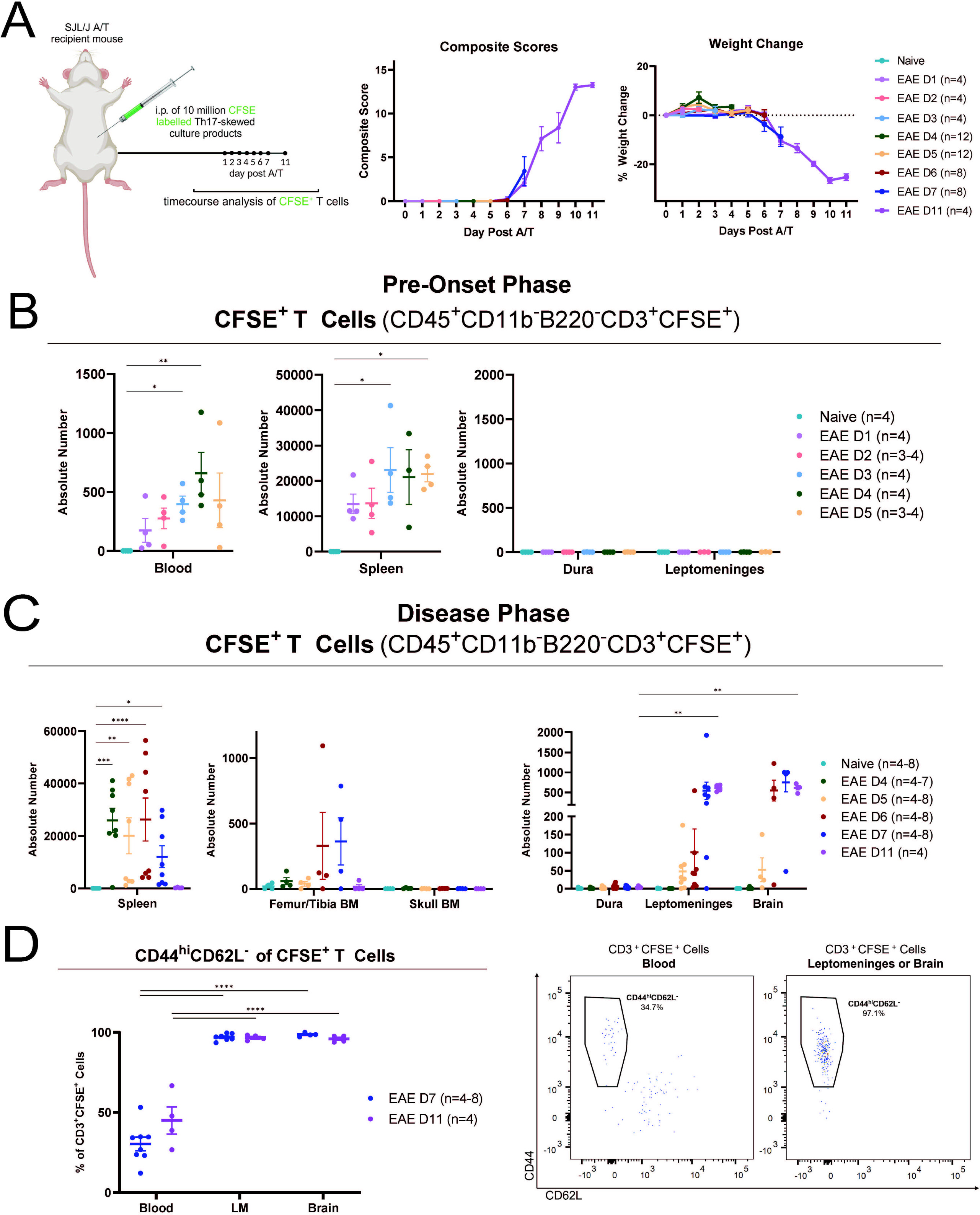
Adoptively transferred encephalitogenic T cells infiltrate the leptomeninges and brain, but not the dura, during the establishment of EAE. Clinical scores were tracked in SJL/J mice in which EAE was induced by adoptive transfer of 10 million CFSE-labelled Th17-skewed encephalitogenic T cells (A). EAE mice developed clinical disability including paralysis (reflected by increasing clinical score) and experienced weight loss (A). At various timepoints throughout disease, mice were euthanized and the dura, LM, brain, blood, spleen and BM from the skull and femur/tibia were collected and analyzed by flow cytometry (B-D). Results are expressed as the absolute number of cells in the whole tissue or 100μL of blood (B and C) or as a percentage of CD3^+^CFSE^+^ cells (D). The percentage of CFSE^+^ T cells that expressed the phenotype CD44^hi^CD62L^-^ was quantified in the blood, LM and brain at day 7 and 11 of EAE (D). Results in B are data from one experiment and results in C and D are data from two independent experiments. A total of 3 to 8 mice were used per timepoint in both B and C as indicated in graph legends. Error bars indicate mean ± standard error of the mean (SEM). For B (blood and spleen) and C (spleen), a Kruskal-Wallis Test followed by Dunn’s multiple comparison test was conducted to assess statistical significance from naïve. For B and C (CNS) and D, a mixed-effects analysis followed by Dunnett’s multiple comparison test was conducted to assess statistical significance from the dura (B and C) or blood (D) within timepoints. For C (BM), a Two-Way ANOVA followed by Sidak’s multiple comparison test was used to assess statistical significance between the leg and skull BM within timepoints. In all cases, statistical significance was denoted as: *= p<0.05, ** = p<0.005, *** = p<0.0005, **** = p<0.0005, no label = not significant. The two variables compared in the mixed-effects analysis/Two-Way ANOVA were days post adoptive transfer (time) and tissue type (B: dura versus LM, C: femur/tibia BM versus skull BM or dura versus LM and brain, D: blood versus LM and brain).

In addition, we tested whether encephalitogenic T cells infiltrate the proximal skull and/or distal femur/tibia BM over the course of EAE. While a few mice had some CFSE^+^ T cells in the femur/tibia BM, the skull BM, like the dura, remained devoid of CFSE^+^ T cell infiltration at any timepoint (Fig. 1C). Lastly, to determine if brain/LM-infiltrating CFSE^+^ T cells expressed the phenotype of naïve (CD44^low^CD62L^+^) or effector memory (CD44^high^CD62L^-^) T cells (Sallusto et al., 1999, Aruffo et al., 1990, Bradley et al., 1992), we analyzed the expression of the activation marker CD44 and the LN entry receptor CD62L, comparing the blood, LM and brain at day 7 and 11 following adoptive transfer. While we observed both CD44^low^CD62L^+^ and CD44^high^CD62L^-^ CFSE^+^ T cells in the blood, almost all CFSE^+^ T cells in the LM and brain were CD44^high^CD62L^-^ (Fig. 1D).

Collectively, these data support previous findings indicating the dura is not a significant reservoir of encephalitogenic T cells in the initiation of EAE (Merlini et al., 2022). Moreover, because this SJL/J version of EAE uniquely invokes Th17 cell-induced LM immune cell aggregates that are adjacent to regions of lymphocyte-poor sub-pial GM lesions (which is reflective of PMS), our study is the first to unambiguously dissociate the dura as a contributing source of such LM-restricted encephalitogenic T cells.

### The leptomeninges is the main site of accumulation of endogenous T and mature B lymphocytes during EAE independent of age

Since both adoptively transferred and endogenous T cells contribute to LM inflammation in SJL/J EAE (Pikor et al., 2015), we next used flow cytometry (Supp. Fig. 1B) to evaluate endogenous T cells at early onset (day 7), late onset (day 9), peak (day 11) and remission timepoints (day 25) in all brain and meningeal compartments of both young and aged mice (Fig. 2A). There was no difference in the number of CD3^+^ T cells in the dura, LM and brain of aged compared to young mice at any timepoint with the exception of an increase in CD3^+^ T cells in the brain of young mice at peak disease. In both young and aged mice, CD3^+^ T cells decreased by remission timepoints despite their highly disparate clinical scores (Fig. 2B). Given that absolute cell numbers in the LM and brain far surpassed those in the dura during disease, we expressed these numbers as a fold-change of CD3^+^ T cells from naïve numbers, comparing the dura to the LM and brain at each EAE timepoint. We observed that CD3^+^ T cells at peak and remission were significantly increased in the LM and brain of young and aged mice compared to the dura (Fig. 2C). Taken together, these data reveal preferential accumulation of CD3^+^ T cells in the LM and brain rather than the dura during EAE, independent of age.

**Fig. 2:**
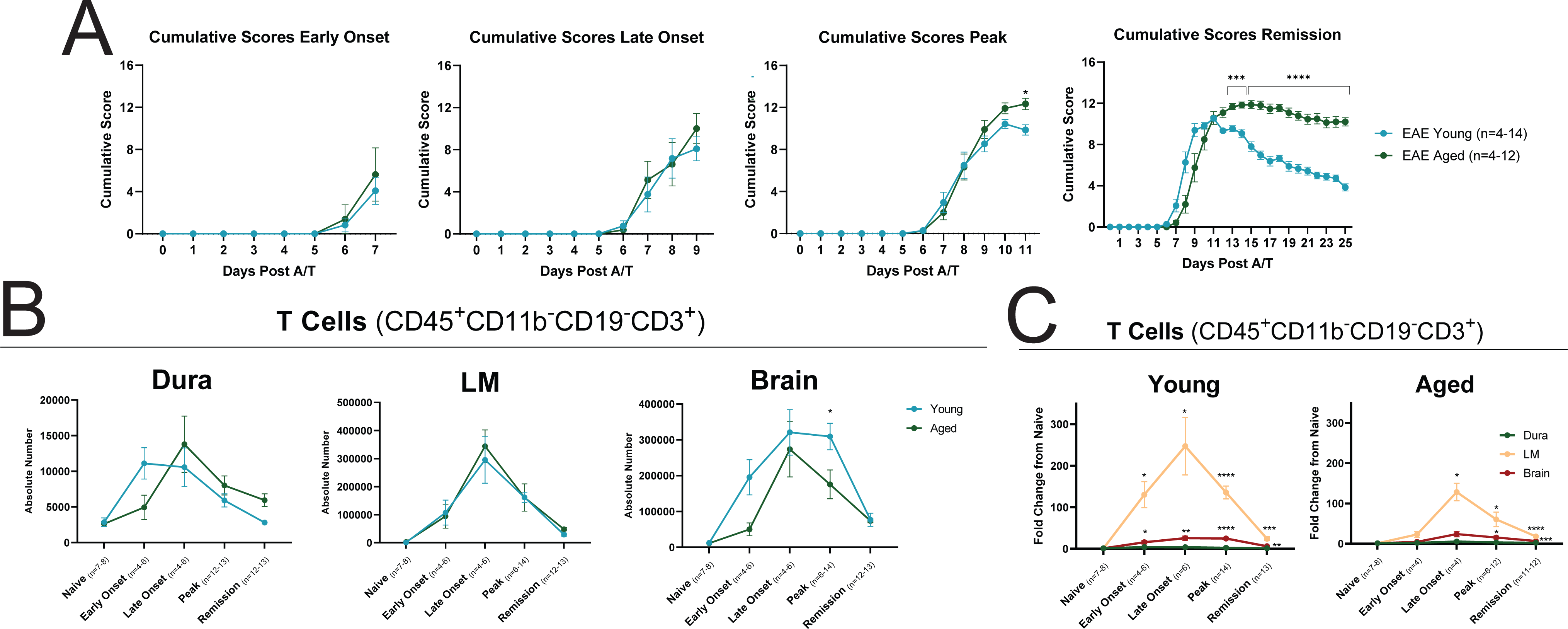
T cells primarily infiltrate the leptomeninges and brain, but not the dura, during EAE independent of age. Clinical scores were tracked in mice in which EAE was induced by adoptive transfer of 10 million Th17-skewed encephalitogenic T cells into young and aged SJL/J mice. Mice developed clinical disability including paralysis (reflected in increased clinical score) (A). At various timepoints throughout disease, mice were euthanized and the dura, LM and brain were collected and analyzed by flow cytometry. Mice were analyzed at naïve (day 0, n=7-8), early onset (day 7, n=4-6), late onset (day 9, n=4-6), peak (day 11, n=6-14) and remission (day 25, n=11-13) timepoints (B to C). Results are data from two repeat experiments. Results are expressed as the absolute number of cells in the whole tissue (B) or as a fold change from naïve mice (C). Error bars indicate mean ± SEM. A Two-Way ANOVA (A and B)/mixed-effects analysis (C) followed by Bonferroni (A), Sidak’s (B) or Dunnett’s (C) multiple comparison test was conducted to assess statistical significance (*= p<0.05, ** = p<0.005, *** = p<0.0005, **** = p<0.0005, no label = not significant). The two variables compared in the mixed-effects model or Two-Way ANOVA were days post adoptive transfer (time) and either age (A and B: young versus aged mice) or tissue type (C: dura versus LM and brain). Of note, statistical significance in graphs plotting immune cell populations as a fold change from naïve in C was evaluated using Dunnett’s by comparing the brain and LM to the dura within each timepoint.

B cells are important contributors to both MS pathology and regulation of neuroinflammation (Wang et al., 2021a). We previously revealed that elimination of LM-resident B cells with anti-CD20 treatment spares the subpial GM from demyelination, synapse loss and evidence of oxidative stress (Wang et al., 2024). To assess the dynamics of mature B cell accumulation in the brain and meninges, we quantified naïve and class-switched B220^high^ B cells, respectively identified as CD19^+^B220^high^CD24^low^IgM^+^IgD^+^ and CD19^+^B220^high^CD24^low^IgM^-^IgD^-^ (Supp. Fig. 1B) in both young and aged mice during the course of EAE (Fig. 2A). In young mice, absolute numbers of naïve B220^high^ B cells increased at onset and peak of EAE and decreased by remission across the brain, LM and dura. Of note, the LM and brain of aged mice harbored significantly fewer naïve B220^high^ naïve B cells at disease peak compared to young mice at the same timepoint (Fig. 3A). A significant fold change in naïve B220^high^ B cells was observed in the LM of both young and aged mice that dwarfed what was observed in the dura. However, in young mice, the brain also showed a significant fold change increase in naïve B220^high^ B cells compared to the dura that was not observed in aged mice (Fig. 3B). Examining class-switched B220^high^ B cells, we observed that these increased in the young and aged dura, LM and brain with EAE but were significantly increased in aged mice in all these tissues at remission (Fig. 3A). Like naïve B220^high^ B cells, the EAE induced fold change in these cells was very modest in the dura (Fig. 3B). Collectively these data show that naïve and class-switched B220^high^ B cells preferentially increase in the LM and brain during the establishment of EAE in young mice, with a preferential accumulation of class-switched rather than naïve B220^high^ B cells in aged mice.

**Fig. 3:**
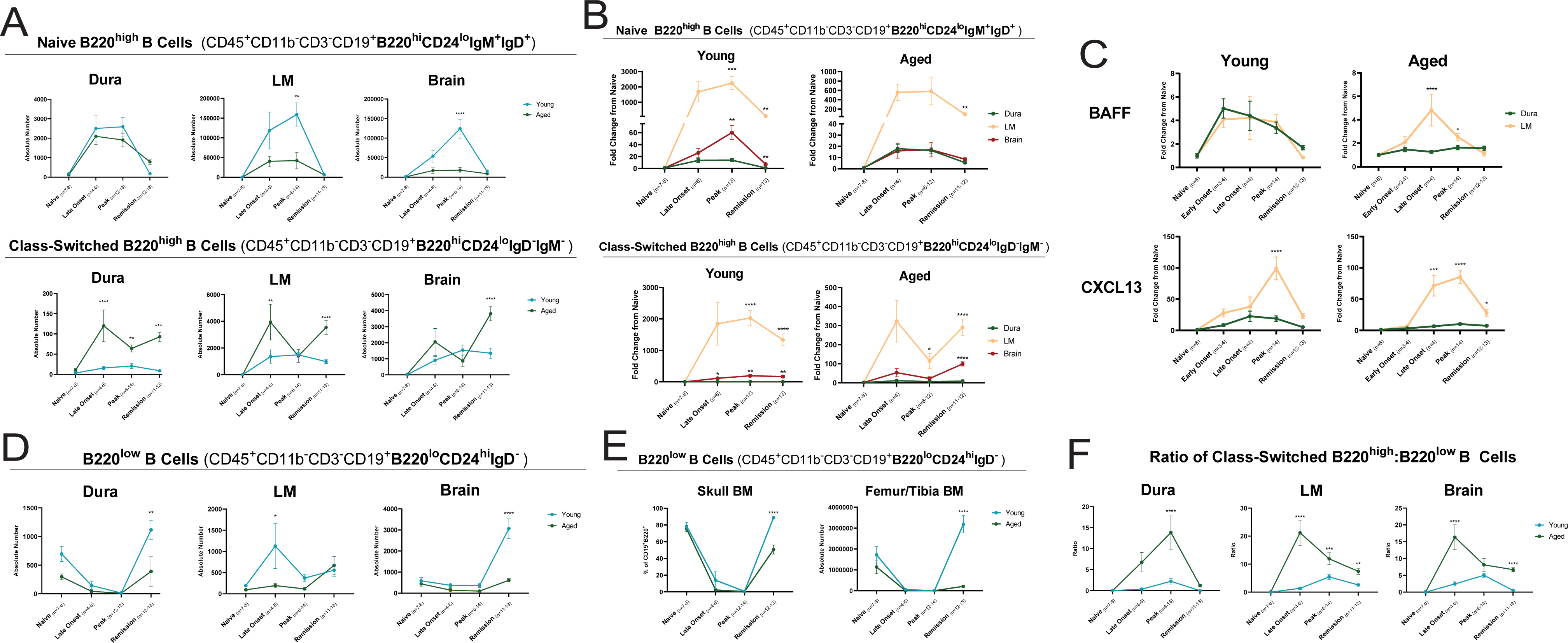
B220_high_ B cells preferentially infiltrate the leptomeninges and brain, but not the dura, of young EAE mice at peak disease, while B220_low_ B cells decrease in the dura and BM but accumulate in the brain of young EAE mice during remission. Clinical scores were tracked in mice in which EAE was induced by adoptive transfer of encephalitogenic T cells into young and aged SJL/J mice (Fig. 2A). At various timepoints throughout disease, mice were euthanized and the dura, LM, brain, skull BM and femur BM were collected and analyzed by flow cytometry. Naïve B220^high^, class-switched B220^high^ and B220^low^ B cells were evaluated at naïve (day 0, n=7-8), late onset (day 9, n=4-6), peak (day 11, n=6-14) and remission (day 25, n=11-13) timepoints. Results are expressed as an absolute number of cells in the whole tissue (A, D, E), a percentage of the parent CD19^+^B220^+^ population (E), as a fold change from naïve mice (B) or as a ratio (F). At the same timepoints, the supernatant of whole tissue dissections of the dura and LM were collected. Concentrations of BAFF and CXCL13 (in pg/mL) were measured in the supernatant using the Ella^TM^ microfluidics platform and results expressed as a fold change from naïve mice (C). Results are data from two repeat experiments. Error bars indicate mean ± SEM. A Two-Way ANOVA followed by Sidak’s multiple comparison test (A, C, D, E and F) or mixed-effects analysis followed by Dunnett’s multiple comparison test (B) was conducted to assess statistical significance (*= p<0.05, ** = p<0.005, *** = p<0.0005, **** = p<0.0005, no label = not significant). The two variables compared in the mixed-effects analysis or Two-Way ANOVA were days post adoptive transfer (time) and either age (A, D, E and F: young versus aged mice) or tissue type (B: dura versus LM and brain, C: dura versus LM). Of note, statistical significance in graphs plotting immune cell populations as a fold change from naïve in B was evaluated using Dunnett’s by comparing the brain and LM to the dura within each timepoint.

We next hypothesized that preferential accumulation of B220^high^ B cells in the LM but not the dura during EAE may be in part explained by differential expression of chemotactic and survival factors. To test this, we measured concentrations of the B cell survival factor B cell activating factor (BAFF) and the B cell attracting chemokine CXCL13 in supernatants from whole tissue extracts of the dura and LM from young and aged mice throughout EAE. Compared to naïve mice, BAFF levels increased in both the LM and dura compartments throughout EAE in young mice. A similar pattern was seen in the LM of aged mice, however unlike young mice, BAFF levels were not elevated in the dura in response to EAE. Like BAFF, CXCL13 also increased in the LM in response to EAE, however there was minimal induction of CXCL13 in the dura at any timepoint in young or aged mice (Fig. 3C). This suggests that an increase in CXCL13 production in the LM may explain the preferential increase in naïve and class-switched B220^high^ B cells in the LM compared to the dura as a result of EAE. The production of CXCL13 within each compartment may be driven by the differential infiltration of Th17 T cells. Indeed, our lab previously showed that interactions between lymphotoxin (LT) αβ on Th17 cells and LTβR on stromal cells in part drove CXCL13 production in the leptomeninges of adoptive transfer SJL/J EAE mice (Pikor et al., 2015). It is therefore possible that greater Th17 cell infiltration into the LM over the dura during EAE drives increased CXCL13 expression and preferential recruitment of B220^high^ B cells to this compartment.

### Identification of anti-CD20 resistant B220_low_ B cells that disappear from the naïve dura and accumulate in the brain during EAE in young but not aged mice

We previously conducted single cell RNA sequencing of the leptomeninges of naïve and EAE SJL/J mice and saw many age-related differences in expression of B cell-associated genes. Of note, compared to aged mice, LM cells derived from young EAE mice showed evidence of upregulation of transcripts associated with developing/immature B cells, including the immunoglobulin superfamily gene expressed in pro- and pre-B cells *Vpreb3* (Zuo et al., 2022). These findings as well as the recent identification of developing B cells in the dura of naïve mice and non-human primates (Brioschi et al., 2021, Schafflick et al., 2021, Wang et al., 2021b) prompted us to examine developing/immature B cells in the dura during EAE. As such, we performed a phenotypic analysis of developing/immature B cells in various tissue compartments during EAE in young and aged mice. Using the BM as a comparator given it is the site of B cell development, we observed two populations of CD45^+^CD11b^-^CD3^-^CD19^+^B220^+^ cells in the dura of young naïve SJL/J mice based on B220 and CD24 expression: B220^high^CD24^low^ and B220^low^CD24^high^ B cells. We observed that CD19^+^B220^high^CD24^low^ B cells were negative for IL7R but expressed major histocompatibility complex II (MHCII) and both IgM and IgD, whereas CD19^+^B220^low^CD24^high^ B cells were variable for IL7R and negative for MHCII and IgD (Supp. Fig. 1B). Therefore, in addition to mature naive B220^high^CD24^low^IgM^+^IgD^+^ B cells, we detected immature/developing B220^low^CD24^high^IgD^-^ B cells in the naïve dura, which for simplicity we term B220^low^ B cells.

We next set out to characterize changes in B220^low^ B cells during EAE. B220^low^ B cells dramatically decreased in the dura at onset and peak EAE in both young and aged mice whilst increasing in the LM of young, but not aged mice at the onset timepoint. In addition, high numbers of B220^low^ B cells appeared in the brain parenchyma of young but not aged EAE mice at remission (Fig. 3D). Moreover, an increased ratio of class-switched B220^high^ to B220^low^ B cells was observed in the LM and brain of aged mice, including at remission (Fig. 3F). Taken together, these data show that B220^low^ B cells virtually disappear from the dura at the same time as their accumulation in the LM and brain during EAE in young mice. However, in aged mice, a deficit in the accumulation of B220^low^ B cells is paired with an increase in class-switched B220^high^ B cells.

Previous studies have suggested that developing B cell populations seed the dura from the overlying skull rather than distal tibial BM (Brioschi et al., 2021). We therefore determined if B220^low^ B cells were altered in these two BM sites in young versus aged mice during EAE. Like the dura, B220^low^ B cells decreased in the skull BM at peak EAE in both young and aged mice. Despite its distance from the CNS, we saw the same decrease in B220^low^ B cells in the femur/tibia BM at peak EAE independent of age. By remission, B220^low^ B cells had rebounded in the femur and skull BM of young mice to levels greater than those seen in naïve mice, but this rebound was not observed in the aged mice who at this timepoint still have chronic EAE (Fig. 3E). It is conceivable that CNS inflammation may trigger a “brain-body” circuit that communicates with the peripheral BM to cause the release of B220^low^ B cells (Jin et al., 2024). Future experiments are required to determine the source of B220^low^ B cells that reach the brain LM and parenchyma (dura versus skull BM versus distal BM).

We recently showed that depletion of B cells with anti-CD20, a therapy that has been largely successful in the treatment of RRMS (Hauser et al., 2017, Hauser et al., 2008, Bar-Or et al., 2008), reduces the size of Th17 cell induced LM immune aggregates and prevents GM demyelination and cortical injury at the peak of disease (Wang et al., 2024). Using an anti-CD20 treatment that effectively depletes B cells in SJL/J mice (Wang et al., 2024), we tested whether B220^low^ B cells are anti-CD20 sensitive (Fig. 4A). Using flow cytometry (Supp. Fig. 1B), we first confirmed that administration of anti-CD20, but not the isotype control, significantly decreased the number of naïve and class switched B220^high^ B cells (Fig. 4B) in all tissues tested in young adoptive transfer EAE mice at remission, whilst sparing CD3^+^ T cells (Supp. Fig. 2). In contrast, the number of B220^low^ B cells in the femur/tibia BM, skull BM and dura were unchanged between isotype and anti-CD20 treated mice at remission. Anti-CD20 treatment resulted in a modest reduction of B220^low^ B cells in the LM and brain, but these were not completely depleted (Fig. 4B). The relative persistence of B220^low^ B cells compared to naïve and class-switched B220^high^ B cells was highly significant in the dura, LM and brain (Fig. 4C).

**Fig. 4:**
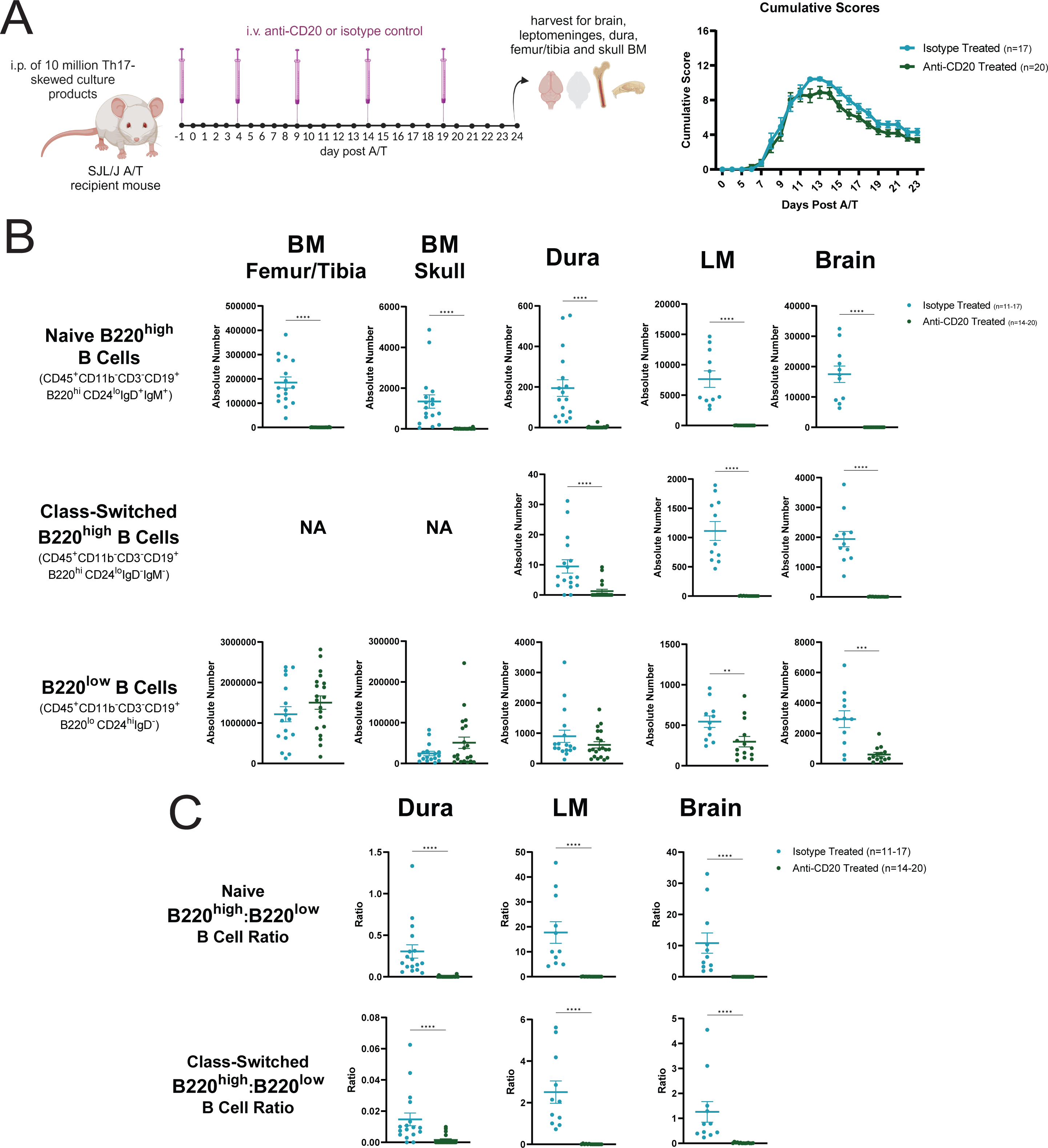
B220_high_, but not B220_low_ B cells are depleted by anti-CD20 therapy in EAE mice. Clinical scores were tracked in mice in which EAE was induced by adoptive transfer of 10 million Th17-skewed encephalitogenic T cells into young SJL/J mice randomized to intravenous treatment with either anti-CD20 monoclonal antibody (n=20) or isotype control antibody (n=17) for on day -1, 4, 9, 14 and 19 of EAE (A). Mice were then euthanized at remission (day 24 of EAE) and the absolute number of naïve B220^high^, class-switched B220^high^ and B220^low^ B cells in the femur/tibia BM, skull BM, dura, LM and brain were analyzed by flow cytometry (B). The ratio of naïve B220^high^ to B220^low^ B cells and class-switched B220^high^ to B220^low^ B cells is also reported (C). The results are data from one experiment. Results are expressed as absolute numbers of whole tissue. Error bars indicate mean ± SEM. A Mann-Whitney Test was conducted to assess statistical significance (*= p<0.05, ** = p<0.005, *** = p<0.0005, **** = p<0.0005, no label = not significant).

Since cortical GM pathology resolves in young mice (Zuo et al., 2022), and B220^low^ B cells persist in the brain at the remission timepoint only in young mice, it is tempting to speculate that B220^low^ B cells contribute towards the prevention of the disease chronicity that we ordinarily observe in aged mice. Moreover, the conspicuous accumulation of class-switched B220^high^ B cells at the expense of B220^low^ B cells in the brain and LM of aged mice, which have severe and persistent cortical GM pathology (Zuo et al., 2022), further supports this hypothesis. Indeed, memory B cells have been implicated in MS pathogenesis (Jelcic et al., 2018, Li et al., 2015).

In summary, in this study we found that T cells and B220^high^ B cells accumulate in the LM and brain during EAE to a much greater extent than the dura, potentially through the induction of CXCL13 within the LM. Moreover, we identified a novel subset of partially anti-CD20 resistant B220^low^ B cells that disappear from the dura and BM yet accumulate in the brain during EAE at a timepoint when young but not aged mice remit from disease, suggesting a potential neuroprotective role for these cells that remains to be investigated.

## Materials and methods

### Mice

Female 3- to 4-week-old SJL/J CD45.1^+^ mice were purchased from Envigo (mouse code: 052). Mice were housed the Division of Comparative Medicine, University of Toronto animal facility under specific pathogen-free conditions, in a closed caging system with a 12-hour light/dark cycle and had free access to a standard irradiated chow diet (Teklad, Envigo, 2918) and acidified water (reverse-osmosis and ultraviolet-sterilized). All experiments were conducted with ethical approval and in compliance with standards for animal care as set out by the Local Animal Care Committee, University of Toronto. Of note, we opted to only use female mice because in the past our laboratory has observed no sex differences in EAE between male versus female mice (Zuo et al., 2022), and male mice present with multiple husbandry issues including penile prolapse resulting in increased need for euthanasia.

### Induction of EAE and clinical evaluation

Donor 6- to 10-week old female SJL/J were immunized with 100 µg of PLP_139-151_ (HSLGKWLGHPDKF; Canpeptide) in an emulsion of incomplete Freund’s adjuvant (BD Difco) and 200 µg of *Mycobacterium tuberculosis* H37 Ra (BD Difco, 231141) via three 100 µL subcutaneous injections on either flank and on the back, for a total injection volume of 300 µL. Nine days following immunization, donors were humanely euthanized by CO_2_ asphyxiation and the spleens and LNs (inguinal, brachial, axillary and cervical) were collected and processed into single cell suspension by mashing through a 70 µm filter under sterile conditions. Cells were then resuspended in complete culture media: RPMI 1640 (Sigma-Aldrich) supplemented with 10% fetal bovine serum (Gibco), 1× penicillin-streptomycin (Sigma-Aldrich), 1× GlutaMAX (Gibco), 10 mM Hepes (Corning), 1 mM sodium pyruvate (Gibco), 100× non-essential amino acids (Gibco) and 1000× β-mercaptoethanol (Gibco). Cells were re-stimulated *ex-vivo* with PLP_139–151_ (10 μg/mL), recombinant mouse IL-23 (10 ng/mL, R&D Systems), anti–IFN-γ (20 μg/mL, Bioceros) and anti–IL-4 (20 μg/mL, Bioceros) for 72 hours at 37°C. 1 × 10^7^ cells from culture were then transferred into young (6- to-10 week old) or aged (8- to 10-month old) female SJL/J recipient mice via intraperitoneal injection on day 0 of EAE. For carboxyfluorescein succinimidyl ester (CFSE) tracking experiments, culture products were first stained at 50 million cells/mL with CFSE (Invitrogen, C34554) in phosphate buffered saline (PBS) at room temperature for 10 minutes with gentle mixing then washed with PBS and transferred into recipient mice via adoptive transfer as described.

For clinical evaluation of disease progression, mice were weighed each day and given a composite clinical score out of a 16-point scoring system that reflects impairments in each limb and tail, as previously described (Galicia et al., 2018, Zuo et al., 2022). Briefly, scores are assigned in 0.5 increments, with each limb given a score out of 3, degree of tail paralysis given a score out of 2 and the righting reflex given a score out of 2. Upon the onset of clinical disease, mice were transferred to a heat pad, provided with wet mash and Napa Nectar (SE Lab Group) and additional hydration was supplied through subcutaneous injections of Lactated Ringer’s Saline. All experiments were pre-approved by the University of Toronto Temerty Faculty of Medicine animal ethics board (protocol #20011363).

### Monoclonal antibody treatment

Young (6- to-10 week old) SJL/J mice were randomized into either treatment or control groups, where the treatment group was administered purified anti-CD20 monoclonal antibody (18B12, mouse IgG2a) generously provided by Roche, while the control group was administered isotype control mouse IgG2a antibodies (BioXCell, BE0085). Both treatment and control groups received 10 mg/kg of antibody dissolved in PBS via five intravenous injections on days -1, 4, 9, 14 and 19 of adoptive transfer EAE. Mice were then humanely euthanized and tissues collected for analysis on day 24 of EAE.

### Preparation for tissue collection

For CFSE tracking experiments, mice were intravenously injected with 3 µg of anti-CD45.1-PE antibody (Thermo Fisher Scientific, 12-0453-82, clone: A20) in 200 µL of PBS three minutes prior to euthanization to preferentially label blood-derived lymphocytes (Ruscher and Hogquist, 2018, Zuo et al., 2022). For all other experiments, mice were immediately humanely euthanized by CO_2_ asphyxiation. Blood was collected into heparin-coated microvette capillary tubes (Sarstedt) via intracardiac bleed and mice were transcardially perfused with 60 mL of ice-cold PBS prior to collection of brain, LM, dura or BM.

### Single-cell isolation from blood

100µL of blood from heparin-coated microvette capillary tubes was then resuspended in 2 mL of red cell lysis buffer (155 mM NH4Cl, 12 mM NaHCO3, and 0.1 mM EDTA) for 4 min on ice. Red blood cell lysis was stopped with the addition of 1 mL of PBS and samples were centrifuged and resuspended in PBS prior to downstream applications.

### Single-cell isolation from LM and dural dissections

The skin overlying the skull was removed and mice were decapitated to separate the skull from the spinal column. The skull cap was then carefully separated from the brain by making incisions from the occipital to the frontal bone to the using fine surgical scissors. The skull cap and brain were each placed into separate petri dishes with 1 mL of PBS. The dura mater was carefully scored from the skull cap and the LM were carefully peeled from the brain under a dissection microscope. Briefly, the dura mater was scored off the skull cap using forceps to release attachment points between the dura and the skull cap whereas the LM were removed from the brainstem, cerebellum, ventricles, hypothalamus, olfactory bulbs and cortex. The dura and LM were then placed into 200 μL and 400 μL of digestion buffer respectively [RPMI 1640 (Sigma-Aldrich) supplemented with 2% fetal bovine serum (Gibco), 1 mM Hepes (Corning), 1× penicillin/streptomycin (Gibco), and 1× GlutaMAX (Gibco)]. After 30 minutes to an hour of sitting in digestion buffer, 200μl of supernatant was removed from the digestion buffer of the dura or the LM and stored at -20°C to - 80°C for downstream protein analysis. Sufficient digestion buffer was then added to top the dura up to 1 mL and the LM up to 500 μL, then digestion enzymes DNase I (dura: 0.5 mg/mL, LM: 60 μg/mL, Sigma-Aldrich) and collagenase P (1 mg/ml, Roche) were added. Tubes were mixed by gentle vortex and digestion allowed to proceed for 15 min at 37°C with gentle agitation. The digestion was stopped with 1 mL of PBS and samples were filtered through 35-μm mesh strainer caps into 5-ml FACS tubes prior to downstream applications.

### Single-cell isolation from brain tissue

Following removal of the LM, the remaining brain tissue was mashed through 70-μm mesh filters into 5 mL of digestion buffer in a 6-well plate. Digestion enzymes DNase I (60 μg/mL, Sigma-Aldrich) and collagenase P (1 mg/ml, Roche) were then added prior to digestion for 30 min at 37°C with gentle agitation. Following digestion, brain homogenates were dissociated by gentle pipetting through the same 70-μm mesh filters. Brain pellets were then resuspended in 30% Percoll (Cytiva) and centrifuged at 2000 rpm for 20 minutes with the brake off to separate lymphocytes from fat and myelin debris. Single cell suspensions of brain were then washed with PBS prior to use for downstream applications.

### Single-cell isolation from skull and leg bone marrow

Following removal of the dura, the skull cap was cleaned of muscle and then cut into small pieces so that pockets of BM were exposed. Skull pieces were then placed into a punctured 0.6mL PCR tube placed within a larger 1.5mL microtube (Sigma). For isolation of the leg BM, the skin of the mouse’s leg was removed and the leg was dislocated from the hip joint. Muscle tissue was cleaned off the femur and tibia and the ends of each bone were cut so as to expose pockets of BM. The femur and tibia were then also placed into a punctured 0.6mL PCR tube placed within a larger 1.5mL microtube. Skull and leg samples were then pulse spun within a microcentrifuge such that the BM from each sample exited the bone and pelleted at the bottom of the 1.5 mL microtube. Samples were washed with 1 mL of PBS and filtered through 35-μm mesh strainer caps into 5-ml FACS tubes prior to downstream applications.

### Flow cytometry

Single cell suspensions were first stained with a 1:1000 dilution of Live/Dead Fixable Aqua (Invitrogen, L34965) for 30 minutes on ice. Surface markers were then stained in 2% fetal bovine serum in PBS (FACS) in the presence of anti-mouse CD16/CD32 Fc block (BioXCell, BE0307) at a concentration of 1/100 for 30 minutes on ice (Supp. Table 1). Where a biotin-conjugated antibody was used in surface staining, samples were subsequently stained with a streptavidin-conjugated fluorophore in 2% fetal bovine serum in PBS. Samples were then fixed and permeabilized in Foxp3/Transcription Factor Staining Buffer (eBioscience, 00-5523-00) fixation/permeabilization diluent for 30 minutes on ice before transferring to FACS buffer overnight. Samples were then mixed at known volumes of FACS with Cell Bright Plus Absolute Counting Beads (Invitrogen, C36995) in order to allow for back calculations of cell numbers. Samples were acquired on a BD Fortessa X-20 or BD FACSymphony A3 using the FACSDiva software and flow cytometry data were analyzed using FlowJo v10.9.0.

### Quantification of proteins from dural and LM supernatant using Ella_TM_ platform

BAFF and CXCL13 concentrations in dura and LM supernatant samples was quantified using a customized mouse cartridge for the Simple Plex Ella^TM^ microfluidics platform (Protein Simple, CA, USA). Samples were thawed and plated neat or diluted at a ratio of 1:2 in manufacturer-supplied diluent. 50 µL of each neat/diluted sample was loaded on the cartridge, which was then placed on the Ella^TM^ instrument. All samples were run in triplicate. Samples were acquired on the Ella^TM^ using factory-generated standard curves corresponding to each cartridge lot. The SimplePlex Explorer (Bio-Techne, v4.1.0.22) was used to retrieve and analyze assay results. The lower limit of quantification (LOQ) and upper LOQ was 10.8 pg/mL and 16478 pg/mL for BAFF and 2.62 pg/mL and 4000 pg/mL for CXCL13.

### Statistical analysis

Unless otherwise stated, all statistical tests were conducted using GraphPad Prism v9.0 and only p-values less than 0.05 were considered significant. The specific statistical tests used in each experiment are indicated in the figure legend text. All quantification data first underwent a Shapiro-Wilk normality test. Flow cytometry data was analyzed using FlowJo v10.9.0. Absolute cell numbers were derived from flow cytometry by recording the number of events in the gate representing the population of interest as well as the number of events recorded in the gate representing counting beads, which were added at a known volume and concentration and allowed for back-calculation of cell numbers as described in the user guide for Cell Bright Plus Absolute Counting Beads (Invitrogen, C36995). Statistical analysis for flow cytometry data was conducted using the following tests without assuming normality: Mann-Whitney test for comparison between two unpaired groups or a Kruskal-Wallis test for comparison between more than two unpaired groups followed by Dunn’s multiple comparison test. Statistical analysis for flow cytometry data comparing two variables was done using either a Two-Way ANOVA or a mixed effects model (if there were missing values in paired data) followed by the following post-hoc tests: Bonferroni’s multiple comparison test for dependent comparisons, Sidak’s multiple comparison test for independent comparisons and Dunnett’s multiple comparison test for independent comparisons to a control group (in this case, the dura).

## Online supplemental material

#Supplementary table 1 lists antibodies used in flow cytometry. Supplementary figure 1 shows flow cytometry gating strategies as pertaining to immune cell populations in figures 1 to 4 in the main body of the text. Supplementary figure 2 quantifies T cell numbers in the BM, dura, LM and brain of anti-CD20 and isotype treated EAE mice at remission.

**Supp. Table 1: Antibodies used in flow cytometry.**

**Supp. Fig. 1: Flow gating strategies.** Flow cytometry gating strategies as pertaining to (A) Fig. 1, (B) Fig. 2 to 4 and Supp. Fig. 2. In A, gating is shown on a representative spleen sample but is applicable to gating in the BM, dura, LM and brain. The same gating is used for blood samples with the exception that double positive CD45 and CD45 i.v. fraction was used for subsequent analysis. In B, two populations of CD45^+^CD11b^-^CD3^-^CD19^+^B220^+^ cells were identified in the naïve dura and femur/tibia BM of a representative SJL/J mouse: mature B220^high^CD24^low^IgM^+^IgD^+^B cells (blue) and immature/developing B220^low^CD24^high^IgD^-^ B cells (red). Further analysis of the surface expression of MHCII and IL7R was analyzed among populations of naïve B220^high^ (blue) and B220^low^ (red) B cells (B). Gating for class-switched B220^high^CD24^low^IgM^-^IgD^-^ B cells (purple) is additionally shown.

**Supp. Fig. 2: T cells numbers are unchanged by anti-CD20 therapy in EAE mice.** Clinical scores were tracked in mice in which EAE was induced by adoptive transfer of 10 million Th17- skewed encephalitogenic T cells into young SJL/J mice randomized to intravenous treatment with either anti-CD20 monoclonal antibody (n=20) or isotype control antibody (n=17) for on day -1, 4, 9, 14 and 19 of EAE (Fig. 5A). Mice were then euthanized at remission (day 24 of EAE) and the absolute number of CD3^+^ T cells in the femur/tibia BM, skull BM, dura, LM and brain were analyzed by flow cytometry. Results are data from one experiment. Results are expressed as absolute numbers of whole tissue. Error bars indicate mean ± SEM. A Mann-Whitney Test was conducted to assess statistical significance (*= p<0.05, ** = p<0.005, *** = p<0.0005, **** = p<0.0005, no label = not significant).

## Abbreviations

BAFF: B cell activating factor
BM: Bone marrow
CNS: Central nervous system
CSF: Cerebrospinal fluid
EAE: Experimental autoimmune encephalomyelitis
GM: Grey matter
LM: Leptomeninges
LOQ: Limit of quantification
MHCII: Major histocompatibility complex II
MS: Multiple sclerosis
PC: Plasma cell
PLP: Proteolipid protein
RRMS: Relapse remitting multiple sclerosis
WM: White mater

## Acknowledgements

We would like the acknowledge the expertise and assistance of Dr. Nathalie Simard in the University of Toronto Faculty of Medicine flow cytometry facility as well as the staff of the Division of Comparative Medicine at the University of Toronto Faculty of Medicine for their animal husbandry.

## Author contributions

AF and JG conceived the study and wrote the original draft of the manuscript; AF, MZ, AAW designed experiments; AF, MZ, AAW, KCJ and MAN performed experiments and edited the manuscript; AF analyzed data; KCJ assisted with statistical analysis; LAW oversaw the purchasing of mice and reagents; EvP and CK supplied the anti-CD20 monoclonal antibody.

## Data and materials availability

All data are available in the main text, in the supplementary materials, or upon request.

