## Supplementary Table 1 for "Dynamic alterations of dural and bone marrow B cells in an animal model of progressive multiple sclerosis"

| Target | Conjugation | Clone | Manufacturer | Catalogue # | Dilution |
|---|---|---|---|---|---|
| B220 | BV711 | RA3-6B2 | BioLegend | 103255 | 1/200 |
| B220 | R718 | RA3-6B2 | BD Biosciences | 567381 | 1/200 |
| CD11b | BUV496 | M1/70 | Thermo Fisher Scientific | 364-0112-82 | 1/200 |
| CD11b | BUV737 | M1/70 | Thermo Fisher Scientific | 367-0112-82 | 1/500 |
| CD11b | BV605 | M1/70 | BioLegend | 101257 | 1/200 |
| CD11b | FITC | M1/70 | Thermo Fisher Scientific | 11-0112-85 | 1/200 |
| CD11c | PE-Cy7 | N418 | Thermo Fisher Scientific | 25-0114-82 | 1/200 |
| CD127 (IL7R) | Biotin | A7R34 | Thermo Fisher Scientific | 13-1271-82 | 1/200 |
| CD127 (IL7R) | PE | A7R34 | Thermo Fisher Scientific | 12-1271-82 | 1/200 |
| CD19 | eF450 | 1D3 | Thermo Fisher Scientific | 48-0193-82 | 1/200 |
| CD19 | RB780 | 1D3 | BD Biosciences | 755522 | 1/200 |
| CD24 | BV750 | 30-F1 | BD Biosciences | 747362 | 1/200 |
| CD3 | BV605 | 17A2 | BioLegend | 100237 | 1/200 |
| CD3 | BV711 | 17A2 | BioLegend | 100241 | 1/200 |
| CD44 | BV605 | 1M7 | Thermo Fisher Scientific | 406-0441-82 | 1/200 |
| CD45 | BUV805 | 30-F11 | Thermo Fisher Scientific | 368-0451-82 | 1/500 |
| CD45.1 (i.v.) | PE | A20 | Thermo Fisher Scientific | 12-0453-82 | i.v. |
| CD62L | APC-eF780 | MEL-14 | Thermo Fisher Scientific | 47-0621-82 | 1/200 |
| CD62L | PE-Cy7 | MEL-14 | Thermo Fisher Scientific | 25-0621-81 | 1/200 |
| I-Ab (MHCII) | FITC | KH74 | BioLegend | 115305 | 1/200 |
| IgD | BV605 | 11-26c.2a | BD Biosciences | 563003 | 1/100 |
| IgM | BUV563 | II/41 | BD Biosciences | 749309 | 1/100 |
| - | BV786 | Streptavidin Conjugated | BD Biosciences | 563858 | 1/200 |

Supplementary Table 1
