## Supplementary figures and images for "Dynamic alterations of dural and bone marrow B cells in an animal model of progressive multiple sclerosis"

### Supplementary Figure 1

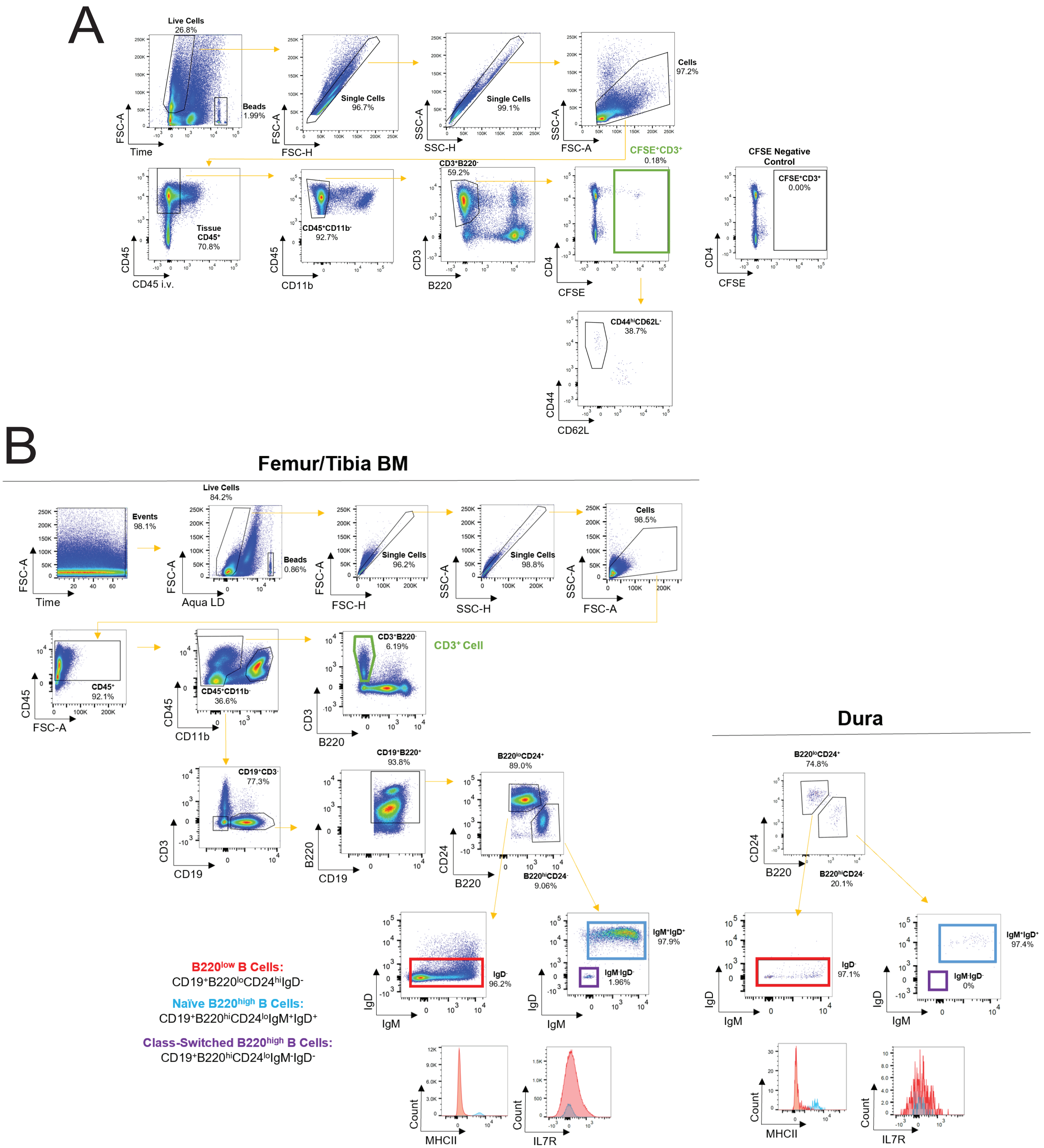


Supplementary Figure 1

### Supplementary Figure 2

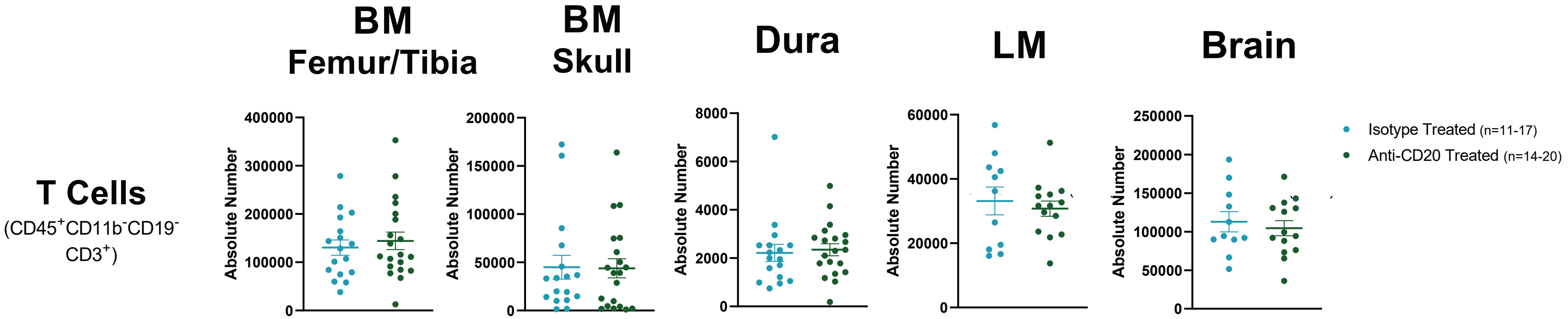


Supplementary Figure 2
